# Masking of a circadian behavior in larval zebrafish involves the thalamo-habenula pathway

**DOI:** 10.1101/097790

**Authors:** Qian Lin, Suresh Jesuthasan

**Affiliations:** NUS Graduate School for Integrative Sciences and Engineering, 28 Medical Drive, National University of Singapore, Singapore 117456.; Lee Kong Chian School of Medicine, Nanyang Technological University, Singapore 636921.; Neural Circuitry and Behavior Laboratory, Institute of Molecular and Cell Biology, Singapore 138673.; Neuroscience and Behavioral Disorders Program, Duke-NUS Graduate Medical School, 8 College Road, Singapore 169857.; Department of Physiology, National University of Singapore, Singapore 117597.

**Keywords:** masking, thalamus, habenula, calcium imaging, diel vertical migration

## Abstract

Light has the ability to disrupt or mask behavior that is normally controlled by the circadian clock. In mammals, masking requires melanopsin-expressing retinal ganglion cells that detect blue light and project to the thalamus. It is not known whether masking is wavelength-dependent in other vertebrates, nor is it clear what higher circuits are involved. Here, we address these questions in zebrafish. We find that diel vertical migration, a circadian behavior in larval zebrafish, is effectively masked by blue, but not by red light. Two-photon calcium imaging reveals that a retino-recipient thalamic nucleus and a downstream structure, the habenula, are tuned to blue light. Lesioning the habenula inhibits light-evoked climbing. These data suggest that a thalamo-habenula pathway may be involved in the ability of blue light to mask circadian behavior.

---

Light has profound effects on animal behavior, independent of visual perception. One effect is mask locomotor activity that is under the control of the circadian clock^1–4^. For example, in diurnal animals such as squirrels, birds and fish, short exposure to light during the night can stimulate movement (positive masking) while exposure to darkness during the day inhibits movement (negative masking). Masking is thought to fine-tune behavior and physiology, such that the animal can respond quickly to changes in the environment. The neural mechanisms underlying masking are unclear^5^. In mammals, masking involves melanopsin-expressing retinal ganglion cells (mRGCs) that are directly sensitive to irradiance, especially of blue light^6,7^. Thus, one approach to identifying neural circuits mediating masking is to examine the projections of mRGCs. By expressing ß-galactosidase^8^ or Cre-recombinase^9^ in the melanopsin locus of mice, or by using viral-mediated tracing^10^, it has been shown that mRGCs innervate distinct targets such as the suprachiasmatic nucleus (SCN), olivary pretectal nucleus as well as thalamic nuclei including the intergeniculate leaflet (IGL) and ventral lateral geniculate nucleus (vLGN). Lesions of the thalamus^11,12^, but not of the SCN^13^, have been reported to affect masking. This indicates an involvement of the thalamus in masking.

Brain regions downstream of the thalamus that contribute to masking are not well defined. One approach to identifying these regions and the underlying mechanism of masking would be to carry out brain imaging using lighting conditions that elicit masking. Here, we do this using larval zebrafish, a system that allows imaging of neural activity across the entire brain at cellular resolution. We first ask whether masking in larval zebrafish^3^ is dependent on wavelength. As a behavioral assay, we use diel vertical migration, which is change in position within a water column. Like many other fish^14,15^, larval zebrafish move to the top of a water column during the day, and to the bottom at night^16^. This behavior can be masked by changes in light^17^, but the wavelength dependency has not been reported. We then image neural activity across the whole brain of larval zebrafish, to identify regions that may mediate masking.

## Results

### Vertical migration is effectively driven by blue light

To characterise vertical migration, larvae were placed independently in custom chambers (Fig. 1a) and exposed to 10 alternating periods of light and darkness, each lasting 1-minute. As previously reported^17^, larval zebrafish moved upwards in the light and downwards in darkness (Fig. 1b). This behavior was seen in fish tested from 5-14 days post fertilization and was induced by a range of intensities (Fig. 1c - e). Larvae showed different climbing speeds under different intensities (Kruskal-Wallis test *χ*^2^ = 6.37, *df* = 2, *p* = 0.0414 for climbing speed; *χ*^2^ = 1.99, *df* = 2, *p* = 0.3693 for diving speed, followed by Mann-Whitney *U* test with Bonferroni correction). Larvae had a slower climbing speed under the low intensity compared to mid intensity (Mann-Whitney *U* = 116, \**p* = 0.006, one tailed, Cohen's *d* = 1.21). The vertical speeds under high and mid intensity were not different (OFF: Mann-Whitney *U* = 91, *p* = 0.28, two tailed, Cohen's *d* = 0.50; ON: Mann-Whitney *U* = 59, *p* = 0.52, two tailed, Cohen's *d* = 0.29). Low intensity of light caused a less robust climbing response, as indicated by the larger confidence interval (Figure 1c), a lower occurrence rate (72 out of 120) and low correlation between the movements of individual fish (Fig. 1e). At the highest irradiance tested, slightly more fish responded in all trials (83 occurrences out of 120). The pair-wise correlation coefficients under mid-intensity light were higher than other light conditions (*χ*^2^ = 9.81, *df* = 2, *p* = 0.0074, Kruskal-Wallis test). Mid vs high: *U* = 2739, *p* = 0.0054; Mid vs low: *U* = 2803, *p* = 0.0022, one tailed Mann-Whitney *U* test with Bonferroni correction. Thus, this condition was used subsequently.

**Figure 1:**
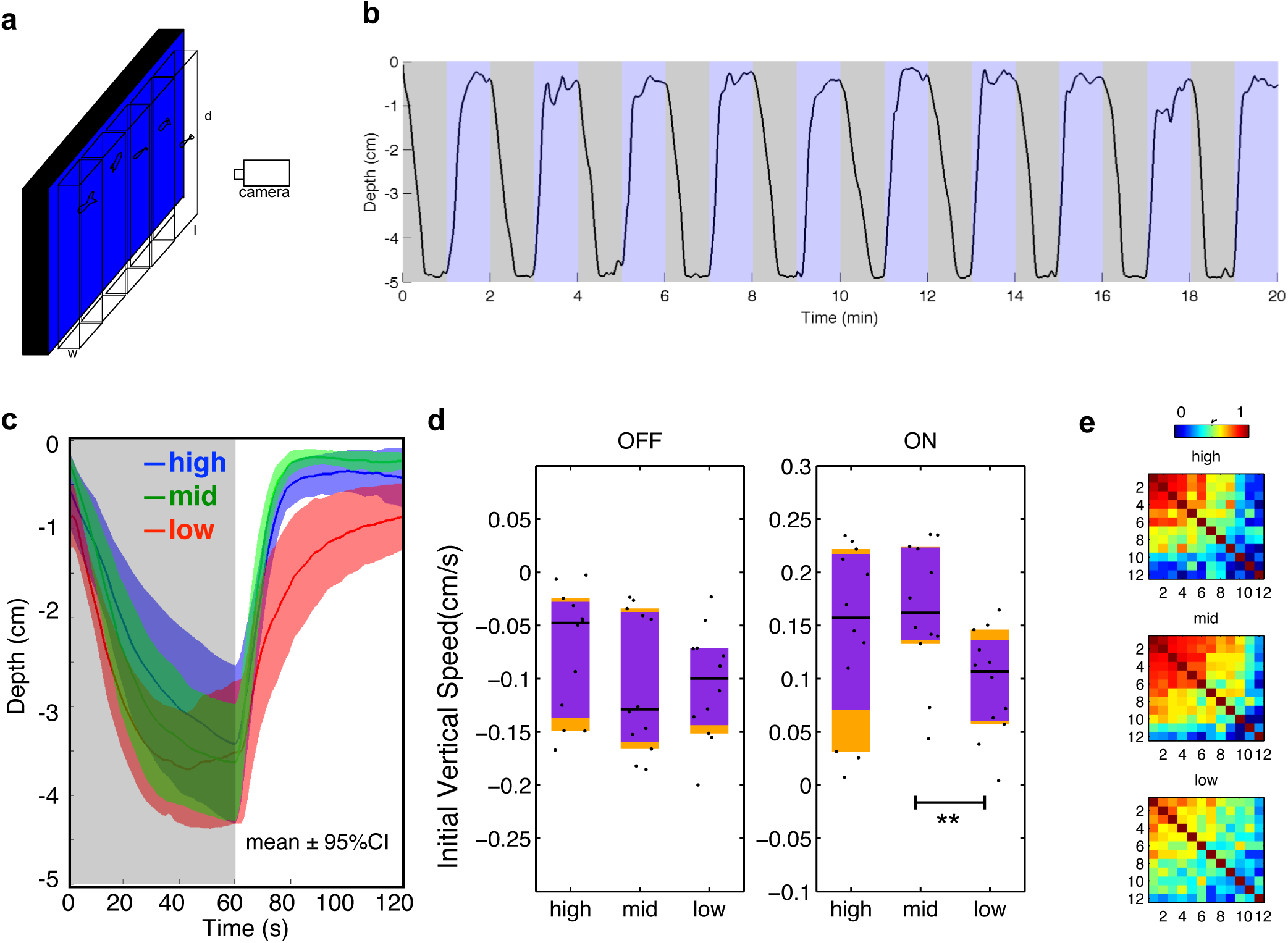
Blue light masks vertical migration. **(a)** Schematic of the assay. Larval zebrafish were placed individually in tanks in front of an LED backlight. **(b)** Response of 1 fish to ten cycles of blue (470 nm) light and darkness. **(c)** Depth of larvae across time under different intensities of blue light, averaged from 10 cycles. n = 12 for each group. Shadows indicate 95% confidence intervals (CIs). Mid-intensity 600 μW/cm^2^, high-level 6000 μW/cm^2^ and low-level is 6 μW/cm^2^. The photon intensity of mid-level is = ~1.56 × 10^15^ cm^−2^s^−1^. **(d)** Comparison of averaged initial 20 s vertical speed under different intensities of blue light. Horizontal black lines indicate median values, black dots show individual speeds, orange blocks show 95% CIs and purple blocks indicate the 25th and 75th percentiles. **(e)** Correlation coefficients of vertical position across time during 10 dark-light cycles under different intensities of light. Correlation matrices are different (mid vs high: *X*^2^ = 6519, df = 66, p < 0.0001; mid vs low: *X*^2^ = 3963, df = 66, p < 0.0001. Jennrich's *X*^2^ test). For n = 3 groups, Bonferroni correction *α* = p / n = 0.05/3 = 0.0167.

Zebrafish have four cones^18^, and are able to detect light across a broad region of the spectrum, from ultraviolet (UV)^19^ to far red^20^. In rodents, UV-sensitive cones contribute to non-image forming response such as circadian photoentrainment and light-induced phase shift^21^. We thus tested whether UV light is able to mask vertical migration of larval zebrafish. Under UV light, at the same photon intensity as blue light, larvae frequently swam up and down, but these movements were not synchronized with periods of light and darkness (Fig. 2a,b). Most fish stayed at the lower portion of the water column during both light onset and offset (Fig. 2b). UV light also led to lower mean vertical speeds (OFF & ON \*\*\**p* < 0.0001, Mann-Whitney *U* test; Fig. 2c) and lower correlation coefficients ( \*\*\**p* < 0.0001; median *r* = 0.11 for UV light and 0.45 for blue light; Fig. 2d). This suggests that masking of vertical migration of larval zebrafish is not caused simply by a change in irradiance, but may be wavelength dependent.

**Figure 2.**
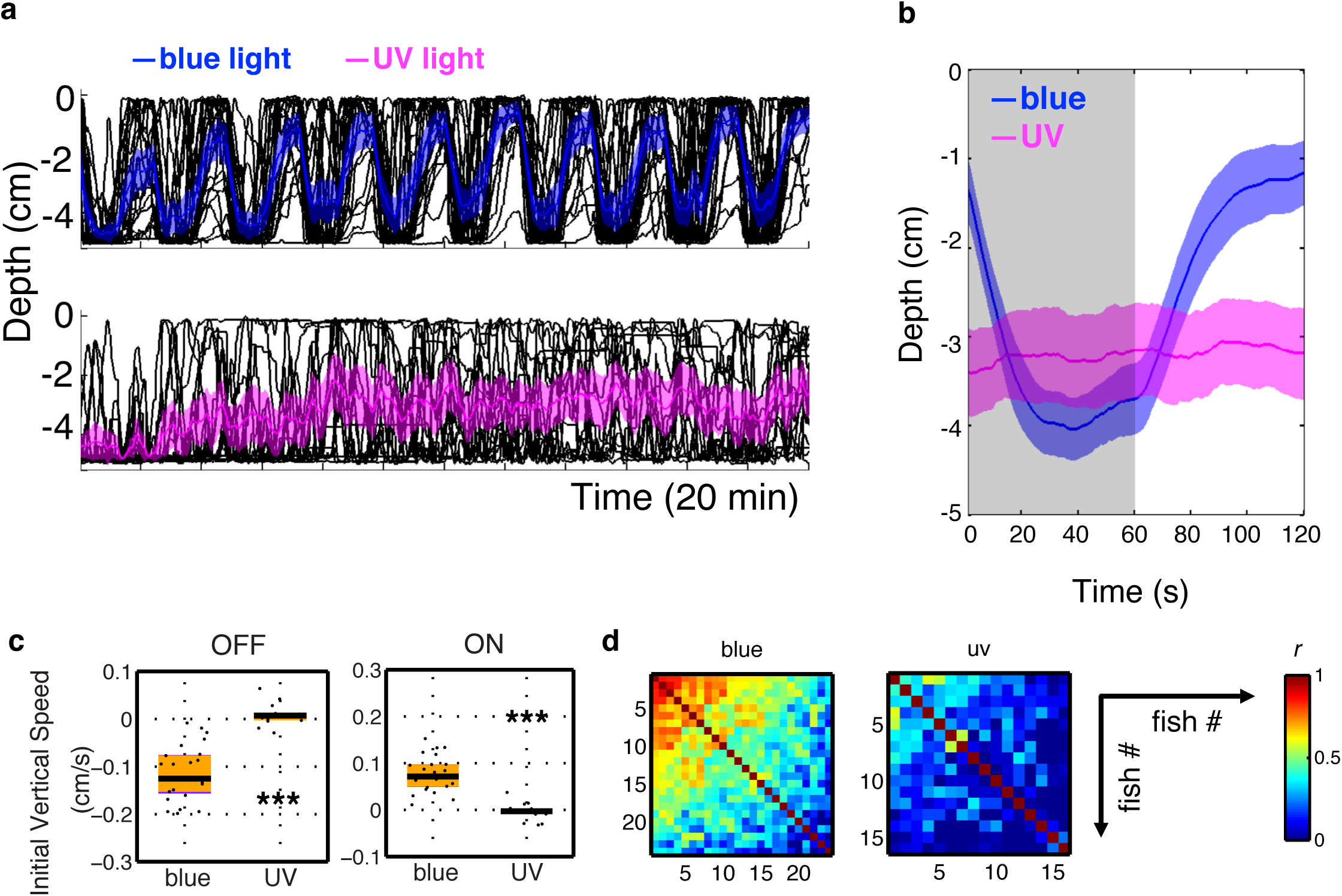
UV light fails to drive vertical migration. (a) Vertical migration of fish under blue (470 nm) and UV (375 nm) light at the same photon intensity, ~1.56 × 10^15^ cm^−2^s^−1^. Black lines represent individual trajectories, and colored traces represent the average of the group. Shadows indicate 95% confidence intervals. n = 24 for the blue group and n = 16 for the UV group. **(b)** Averaged vertical migration from 10 dark-light cycles. **(c)** Initial 20 s diving and climbing speed. Vertical speeds under UV are reduced relative to the speeds under blue light (light OFF: Mann-Whitney *U* = 380, \*\*\**p* < 0.0001, two tailed, Cohen's *d* = 2.82; light ON: Mann-Whitney *U* = 378, \*\*\**p* < 0.0001, two tailed, Cohen's *d* = 2.76). Horizontal black lines indicate median values, black dots show speed of individual fish, purple blocks show the 25th and 75th percentiles and orange blocks show 95% CIs. **(d)** Correlation coefficient of vertical position across time during 10 dark-light cycles. Correlation coefficient is lower under UV light than under blue light (*U* = 3442, *p* < 0.0001, one tailed Mann-Whitney *U* test; median *r* = 0.11 for UV and 0.45 for blue).

To test this further, we examined the effects of red and green light, in comparison to blue. Under the same photon intensity, green light triggered a similar vertical migration as blue light, with similar depth over time (Fig. 3a, b), diving/climbing speeds (two-tailed *p* > 0.7; Fig. 3c), and high correlation coefficient across 10 cycles among individual fish (median *r* = 0.67 for blue, *r* = 0.69 for green; two-tailed *p* = 0.53; Fig. 3d). In contrast, red light failed to drive a full vertical migration (Fig. 3a, b), with a less synchronized up and down movement, causing a lower correlation coefficient among individuals (median *r* = 0.40, \*\**p* < 0.0001; Fig. 3d). Fish also failed to remain at the top of the water column under red light (Fig. 3b), and there was decreased diving/climbing speeds (compared to blue light, diving: *one-tailed *p* = 0.0097; climbing: two-tailed *p* = 0.040; Fig. 3c). In addition, when the same photon intensity of red and blue light were given alternatively, larvae displayed a small but obvious vertical migration, diving upon red light onset and climbing upon blue light onset, leading to different vertical positions (**two-tailed *p* = 0.008; Fig. 3e). Thus, change in red light has distinct effects from change in green and blue light.

**Figure 3.**
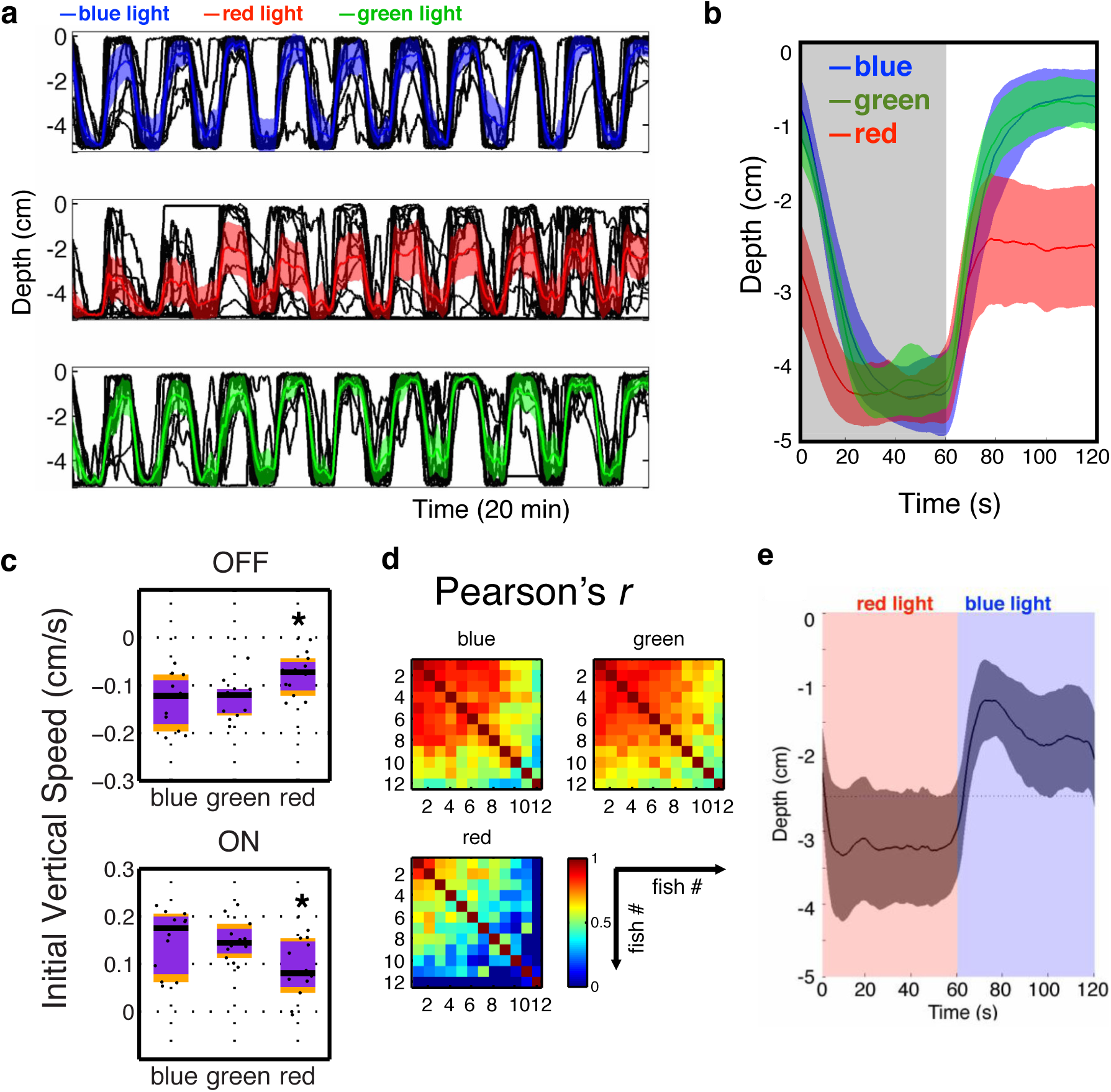
A comparison of vertical migration under blue, green and red light. (a) Vertical migration of larvae exposed to 10 cycles of blue (470 nm; 650 μW/cm^2^), green (525 nm; 580 μW/cm^2^) and red (660 nm; 465 μW/cm^2^) light, which yield the same photon intensity, ~1.56 × 10^15^ cm^−2^s^−1^. Black lines represent individual fish, while colored traces represent the average. Shadows indicate 95% CIs. n = 12 for each group. **(b)** Mean vertical migration from 10 dark-light cycles, averaged from panel a, represented as mean ± 95% CIs. **(c)** Comparisons of initial 20 s vertical swimming speeds. Vertical speeds under blue and green lights are similar (Diving: Mann-Whitney *U* = 66, *p* = 0.76, two tailed, Cohen's *d* = 0.13; climbing: Mann-Whitney *U* = 76, *p* = 0.84, two tailed, Cohen's *d* = 0.0093). Vertical speeds under blue and red light are different (Diving: Mann-Whitney *U* = 113, *p* = 0.0097, one tailed, Cohen's *d* = 1.16; Climbing: Mann-Whitney *U* = 108, *p* = 0.040, two tailed, Cohen's *d* = 0.97). Vertical speeds under green and red light are different (Diving: Mann-Whitney *U* = 114, *p* = 0.0083, one tailed, Cohen's *d* = 1.20; Climbing: Mann-Whitney *U* = 110, *p* = 0.015, one tailed, Cohen's *d* = 1.19). **(d)** Correlation coefficient of vertical position across time during 10 dark-light cycles. Correlation matrices are different (blue vs green: *X*2 = 3288, *df* = 66, *p* < 0.0001; blue vs red: *X*2 = 5534, *df* = 66, *p* < 0.0001. Jennrich's *X*2 test). **(e)** Behavior under alternating red and blue light cycles with the same photon intensity ~1.56 × 10^15^ cm^−2^s^−1^, represented as mean ± 95% CIs. Larvae display vertical migration, with the vertical position under blue light being higher than the vertical position under red light (Mann-Whitney *U* = 118, *p* = 0.008, two tailed, Cohen's *d* = 1.34).

The deficits under red light could be because red light acts through a different light processing pathway that does not drive a full vertical migration response, or because the fish are less sensitive to red light. To investigate the latter possibility, red light of 10-fold higher in intensity was applied, to compensate for the reduced sensitivity. Under this higher intensity of red light, larvae still performed an incomplete vertical migration (Fig. 4a), with decreased depth during light onset (Fig. 4b), a decreased diving speed (\*\**p* = 0.004; Fig. 4c), and lower synchronization among individuals (median Pearson's *r* = 0.15 in contrast to median *r* = 0.45 for blue; *p* < 0.0001; Fig. 4d). Although there was a similar climbing speed upon light onset (Fig. 4c), larval fish under bright red light did not maintain their vertical position during the light-on period (Fig. 4b). This different behavior implies that changes in blue and red illumination are not processed in the same way.

**Figure 4.**
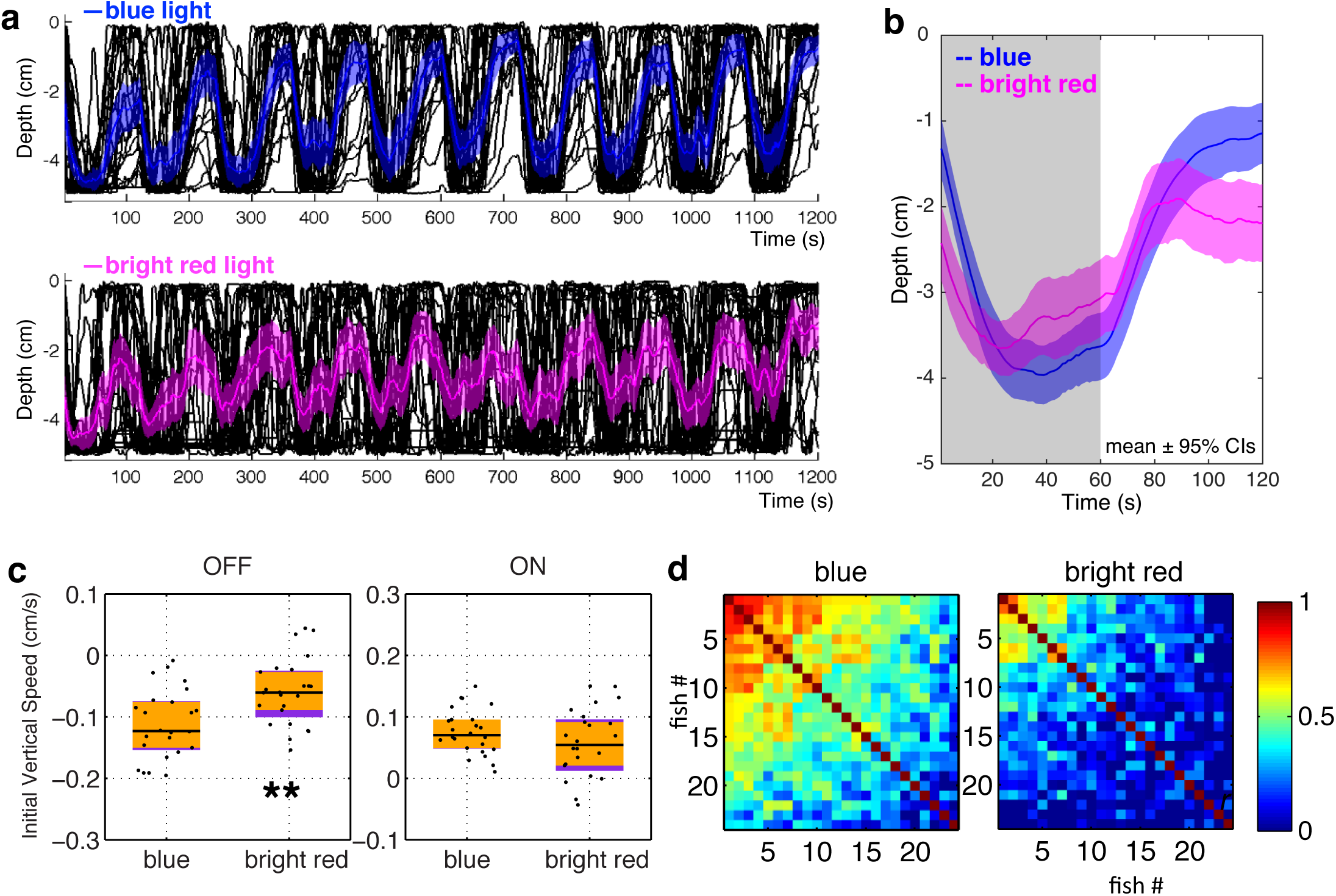
Irradiance change in blue and red have different effects. **(a)** Vertical migration of fish under blue light and red light of ten-fold higher intensity: ~1.56 × 10^15^ cm^−2^s^−1^ for blue light, and ~1.6 × 10^16^ cm^−2^s^−1^ for red light. Black lines represent individual fish, and colored traces represent the average. Shadows indicate 95% CIs. Blue, 470 nm, 650 μW/cm^2^; red, 660 nm, 4650 μW/cm^2^. N = 24 for each group. **(b)** Averaged vertical migration from 10 dark-light cycles, averaged from panel a, represented as mean ± 95% CIs. The curve under red light of high intensity is different from the one under blue towards the end of the light ON period. (c) Initial 20 s vertical swimming speed. Initial diving speeds upon light offset are different (Mann-Whitney *U* = 431, \*\**p* = 0.004, two tailed, Cohen's *d* = 0.95). Initial climbing speeds upon light onset are similar (Mann-Whitney *U* = 345, *p* = 0.24, two tailed, Cohen's *d* = 0.88). Horizontal black lines indicate median values, black dots show speed of individual fish, purple blocks show the 25th and 75th percentiles and orange blocks show 95% CIs. **(d)** Correlation coefficient of vertical position across time during 10 dark-light cycles. Correlation matrices are significantly different (*X*^2^ = 12975, *df* = 276, *p* < 0.0001. Jennrich's *X*^2^ test). Correlation coefficients are lower under red light of high intensity compared to blue light (median *r* = 0.45 for blue, 0.15 for red; *U* = 2803, \*\*\**p* < 0.0001, one tailed Mann-Whitney *U* test). **(e)** The effect of wavelength on the visual-motor response over multiple cycles of light and dark. n = 24 in each group. The trace shows average value, while bars indicate 95% CIs. The black bar indicates period of darkness. The color of the trace represents the wavelength of light preceding the dark phase. Blue and red lights were given in alternation. In all cases, the response to light ON and OFF was stronger in the cycle with blue light.

### Calcium imaging identifies potential correlates of blue sensitivity

The difference between the effects of red and blue light provides a tool to probe the neural mechanism of masking. Specifically, neurons that are activated by changes in blue light, but not by changes in red light, are candidates for components of the circuits that mediate masking; cells that respond similarly could be those that detect visual stimuli in general. To identify these neurons, we performed calcium imaging of fish with broad expression of GCaMP6f^22^, using resonant-scanning two-photon microscopy. Fig. 5 shows the comparison of neural activity evoked by blue and red light. One-minute pulses of red or blue light, alternating with 1-minute darkness, were delivered in a random fashion. Images were processed by principal component analysis (PCA) to reduce dimensionality and then by independent component analysis (ICA) to obtain separate signals^23^. This analysis suggests that there are several responses to increase in irradiance. Blue-specific sustained excitation (Fig. 5c) was detected in the thalamus (Fig. 5a, b; blue pixels; arrowheads). A wavelength-independent transient excitation to onset and offset of light (Fig. 5d, e) was prominent in the tectal neuropil (pink and yellow pixels; Fig. 5a). Activity was also detected in the left habenula (Fig. 5a arrow), including sustained excitation to blue light and transient excitation to darkness.

**Figure 5.**
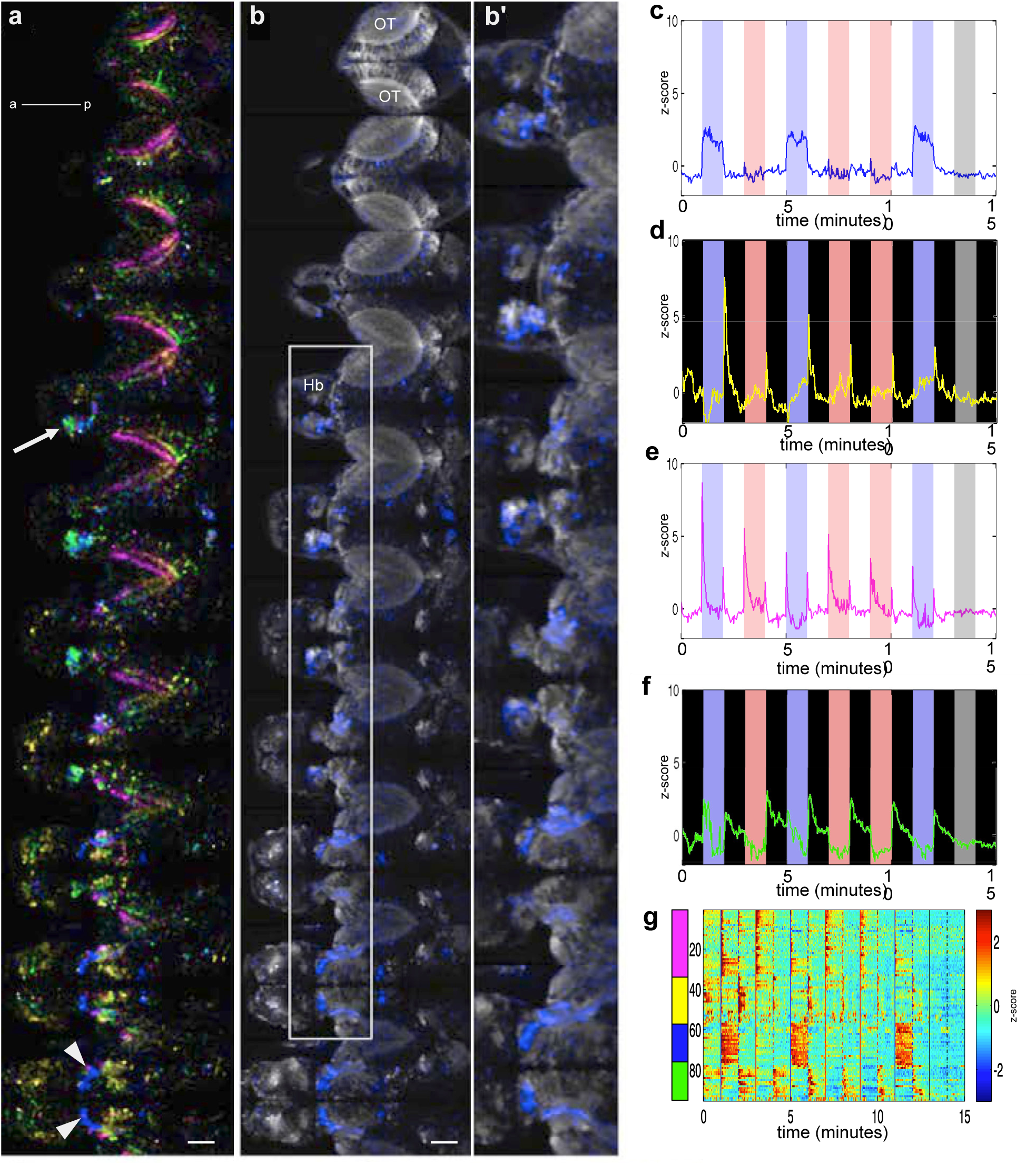
Overview of blue versus red light-evoked activity in the brain. **(a)** Optical sections in a 6 dpf *elavl3:GCaMP6f* fish, going from dorsal to ventral. Each plane is separated by 10 μm. The area imaged at each plane is 443.47 μm × 222.73 μm. **(b)** Distribution of light-evoked activity in one fish, generated from ICA spatial maps. The white arrowheads indicate the thalamus, while the arrow indicates the habenula. **(b')** A magnified view of the area boxed in panel **b**. **(c-f)** Different classes of responses, obtained by PCA followed by ICA. The same colours are used in panel **b**. The blue and red bars indicate the periods that blue or red light was delivered. **(g)** Light-evoked activity in segmented ROIs, which were derived from ICA spatial maps. OT: optic tectum; Hb: habenula; a = anterior; p: posterior. Gamma of 0.65 was applied to panels a and b. Scale bar = 50 μm.

Given that temporal signals derived from ICA were averaged by weights of each pixel, it is possible that real fluorescence signals might be different. To test this, ROIs were segmented from the ICA spatial maps by thresholding. The activities of individual ROIs were then averaged and presented as z-scores (Fig. 5g). Signals from segmented ROIs, such as the sustained excitation to blue light, were similar to their corresponding ICA temporal signals, indicating that ICA signals accurately reflect neural activity reported by the calcium indicator.

To assess reproducibility, the neural responses of 7 fish were collected and integrated (Fig. 6; see Methods). Averaged ICA spatial maps show that the optic tectum has wavelength-independent phasic excitation in response to onset or offset of light (Fig. 6c, f). Darkness caused transient excitation in the optic tectum and the habenula (Figure 6b, e). A sustained response to blue light was seen in the habenula and the thalamus (Fig. 6a, d).

**Figure 6.**
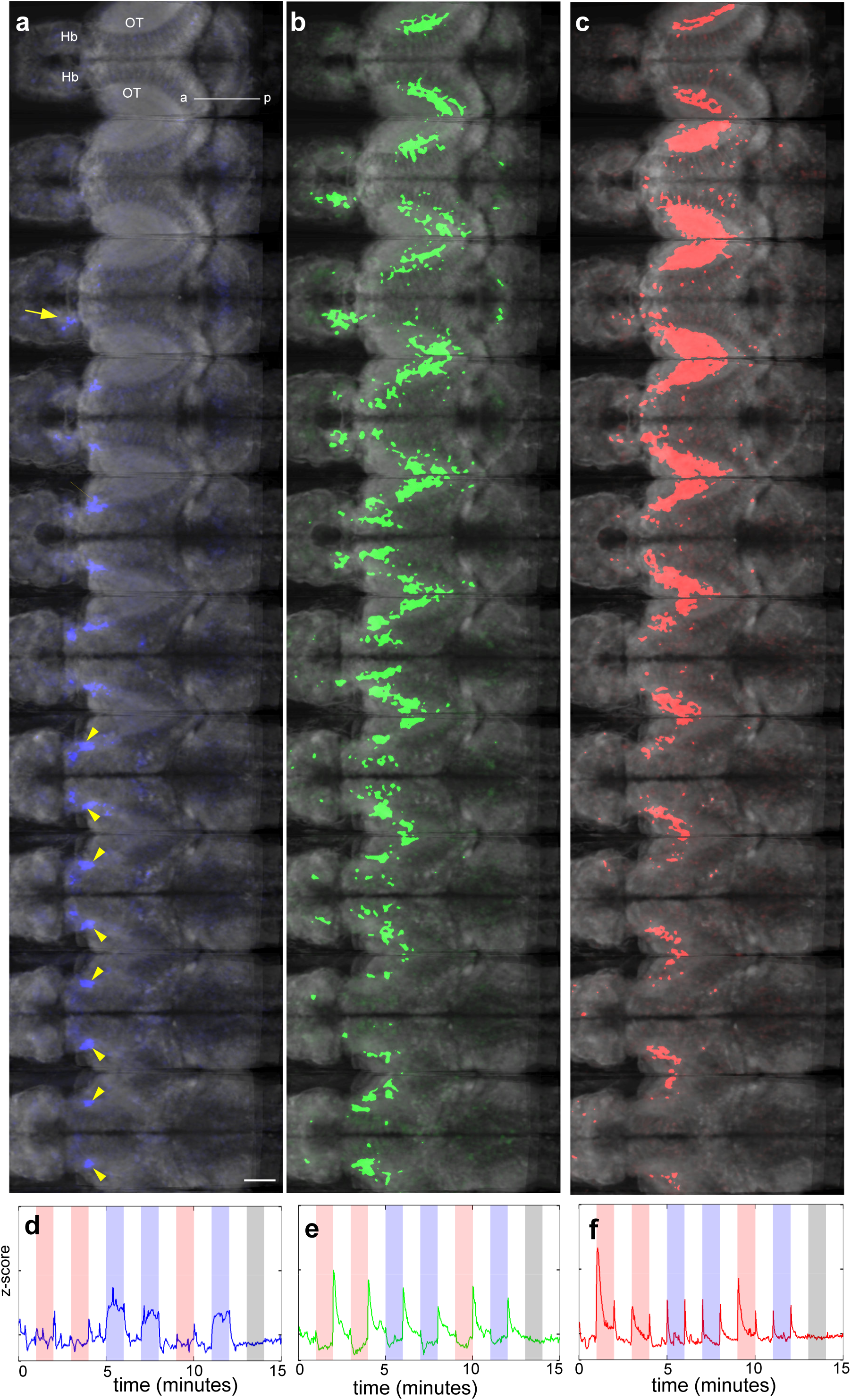
Averaged response of the brain to irradiance change. (**a-c**) Regions with sustained response to blue light **(a)**, to offset of light **(b)** and to onset and offset of both blue and red light **(c)**. The corresponding activity traces are shown in panels **d**, **e** and **f**. These data are compiled from 7 fish.

### Lesioning the habenula affects masking

The difference in habenula response to blue relative to red light suggests that the habenula may be involved in the effect of blue light on vertical migration. To test this, we recorded behaviour after lesioning the habenula. The zebrafish habenula has two major subdomains, dorsal and ventral, and dorsal subdomains are asymmetric. Imaging of the habenula with a galvano scanner, which provides a better signal-to-noise ratio, has shown that blue light evokes activity strongly in the dorsal left neuropil and also bilaterally in the ventral habenula^24^. We lesioned different regions of the habenula, using a femtosecond laser, to test their involvement in masking.

Fish with the dorsal left neuropil lesioned (Fig. 7a, Movie 1) displayed a decreased climbing speed in the presence of light (*U* = 551, \**p* = 0.020, Cohen's *d* = 0.68), but diving speeds during darkness were not statistically different (*U* = 329, *p* = 0.83, Cohen's *d* = 0.020) (Fig. 7b, c). Consistent with this, lesioned fish took a longer time to climb half of the water column during the light phase, while the time to dive half of the water column during the dark phase were not different (Fig. 7d). There was also a slight decrease in the correlation of movement (Fig. 7e; median *r* = 0.45 for the lesion group and 0.51 for the control group; *U* = 63217, \*\*\**p* < 0.0001. N = 29 for control group, 28 for lesion group.). Lesioning the right dorsal neuropil (Fig. 7f) had no significant effects on the speed of diving during light OFF or climbing during light ON (Fig. 7g-I; OFF: *U* = 176, *p* = 0.268, two tailed, Cohen's *d* = 0.371; ON: *U* = 271, *p* = 0.208, Cohen's *d* = 0.428), but there was a difference in the correlation of movement (Fig. 7j; Median r = 0.41 for lesion group, 0.58 for control group. *U* = 29200, \*\*\**p* < 0.0001. N = 21 for each group). When the ventral neuropil was lesioned (Fig. 7k; Movie 2), diving and climbing speeds differed significantly from control group (OFF: *U* = 219, \*\**p* = 0.004, Cohen's *d* = 0.819; ON: *U* = 664, \*\*\**p* < 0.0001, Cohen's *d* = 1.45). Swimming in the lesioned group was also less synchronized compared to the control group (Fig. 7o; median *r* = 0.15 for the lesion group and 0.51 for the control group; *U* = 23885, \*\*\**p* < 0.0001; N = 29 for control group, 27 for lesion group). These results are consistent with the hypothesis that the habenula is involved in the ability of blue light to mask diel vertical migration.

**Figure 7.**
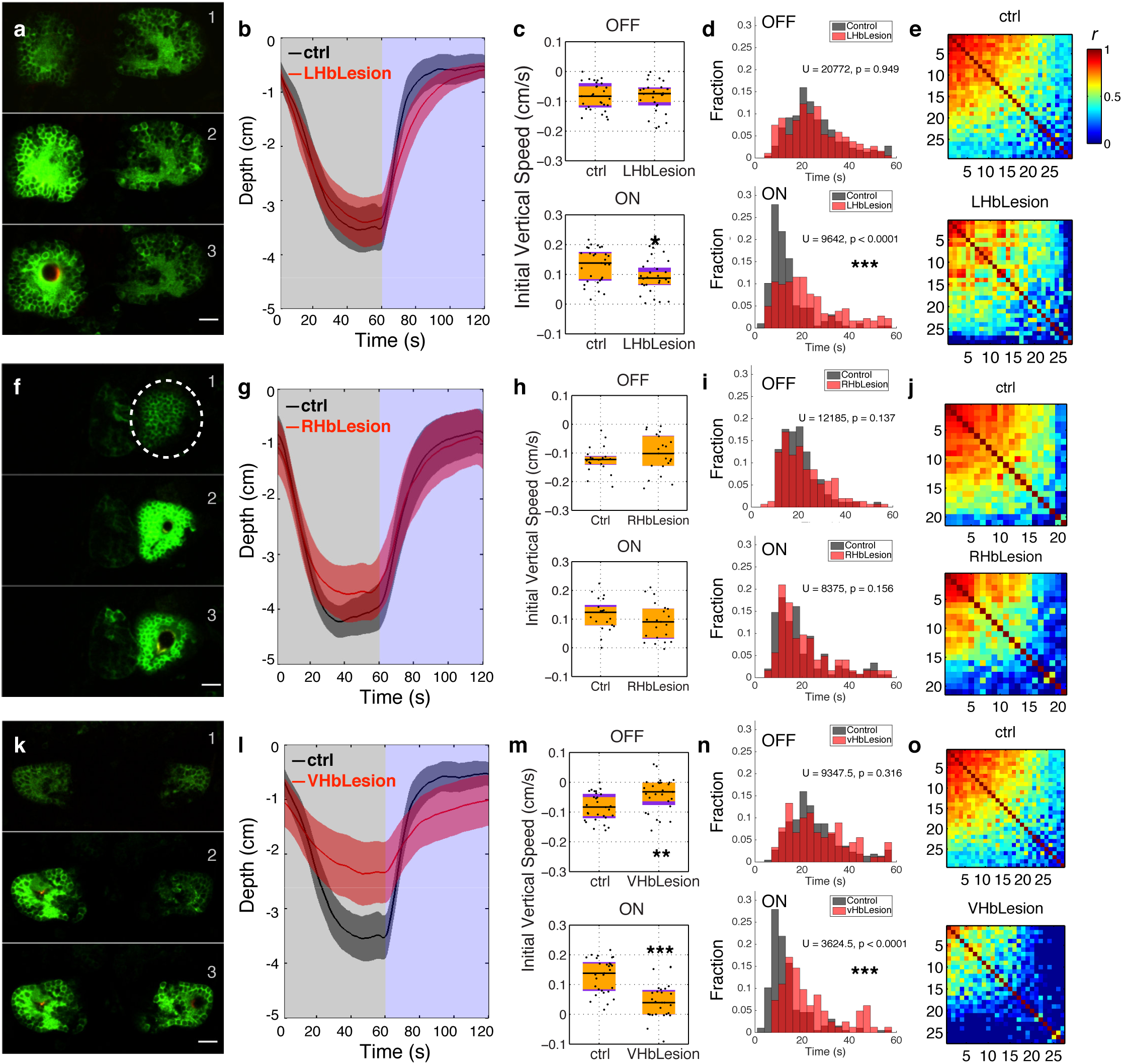
Lesion of the habenula inhibits masking of vertical migration. Effects of lesioning the left dorsal (**a-e**; N = 29 for control group, 28 for lesion group), right dorsal (**f-j**; N = 21 for each group) and ventral (**k-o**; N = 29 for control group, 27 for lesion group) habenula. **(a, f, k)** Stills from a time-lapse recording taken during the lesioning process, showing the habenula before (top), during (middle) and immediately after (bottom) lesioning. Lesioning leads to the formation of a cavity and extended rise in GCaMP3 fluorescence. **(b, g, l)** Average position of fish during light ON and OFF. Shadows represent 95% CIs. **(c, h, m)** Initial vertical speed during light ON and OFF. Horizontal black lines indicate median values, black dots show individual fish, purple blocks show the 25th and 75th percentiles and orange blocks show 95% CIs. **(d, i, n)** Time taken to migrate half-way up or down the water column. The bars show the percentage of fish that reach this mark within the indicated time. **(e, j, o)** Pair-wise correlation between the movement of individual fish.

## Discussion

In this manuscript, we provide evidence that blue light drives vertical migration in larval zebrafish, and also stimulates a thalamic nucleus and the habenula. By lesioning, we show that the habenula influences the effect of blue light on vertical migration. As the thalamus directly innervates the habenula and appears to be the only pathway mediating the response of the habenula to light^24^, these results indicate that a thalamo-habenula projection is involved in the ability of blue light to mask a circadian behavior. Given that the habenula is a regulator of neuromodulators, these results imply that blue light influences motor activity via the release of neuromodulators.

The thalamic nucleus involved here is likely to be the nucleus rostrolateralis^25, 26^, a structure that has been defined in fish, but whose homology to nuclei in the mammalian thalamus is undefined. The neuropil of the nucleus rostrolateralis may include the arborization field AF4^24,27^. AF4 is innervated by M3 and M4 retinal ganglion cells^28^, which are both “ON” neurons. Based on the sustained, eye-dependent excitation in the thalamic neuropil in the presence of blue light^24^, it is possible that some M3 or M4 neurons are either intrinsically sensitive to blue light, or receive input from other retinal cells that express melanopsin-related genes^29^.

There are several lines of evidence to suggest that a thalamic nucleus with blue sensitivity and connectivity to the habenula is not a feature that is specific to larval fish. In the frog, electrical recordings have demonstrated a preferential response to blue light in the dorsal thalamus^30^ and retrograde tracing shows that the habenula of amphibians is innervated by the dorsal thalamus^31^. In the mouse, melanopsin expressing retinal ganglion cells innervate the thalamus, and an analysis of the mesoscale connectome^32^ shows that the ventral lateral geniculate nucleus of the mouse thalamus innervates the habenula. In the rat, anterograde tracing suggests that neurons from the thalamus innervates the habenula^33^. In humans, blue light activates the thalamus^34^, which has functional connectivity with the habenula^35^. It is thus possible that a thalamo-habenula projection plays a role in the ability of blue light to mask behavior in vertebrates.

## Methods

### Zebrafish lines

Experiments were carried out in accordance with protocols approved by the Institutional Animal Care and Use Committee. Zebrafish (*Danio rerio*) lines used in this study were: *elavl3:GCaMP6f*^24^, *s1011tGAL4*^36^, *UAS:GCaMP3*^37^, and AB wildtype (http://zfin.org/action/genotype/view/ZDB-GENO-960809-7).

Imaging was carried on larvae in a *nacre*^−/−^ background^38^. Animals were housed in a facility with lights on between 8 am and 10 pm, and were chosen randomly for experiments.

### Climbing assay

Naive larvae (1 - 2 weeks old) were placed individually in a chamber (3.0 cm length × 1.0 cm width × 5.0 cm height), and housed inside an incubator to exclude light. After 3-8 min adaptation to light, larvae were exposed to 10 cycles of alternating light/dark, each consisting of 60 seconds light ON and 60 seconds light OFF. Blue (470 nm peak), green (525 nm peak) or red light (660 nm peak) was provided by a LED backlight (TMS; see http://tms-lite.com/wavelength/ for spectra). Photon intensity, which is the relevant measure for the non-visual system^39^, was calculated using the equation I = P/(A*E). Power, P, was measured using a meter (ThorLabs PM100A) and A is the receptive area of the photodiode power sensor (Thorlabs S120VC). E, the energy of a single photon, was calculated with the equation E = hc /λ, where c is light velocity and h is Planck's constant.

6 fish were tested simultaneously. Videos were recorded at 5 fps, 640 × 480 pixel resolution, using MicroManager^40^ to control a Grasshopper camera (Point Grey Research) equipped with a 12 mm lens and 830 nm longpass filter. Four infrared LED bars (TMS-lite, Model: LBS2-00-080-3-IR850-24V, 850 nm) were used for illumination. The light box was controlled by a power supply analogue (TMS-lite), which was triggered by a microcontroller board (Arduino UNO SMD) and Arduino software. All animals for a given experiment were from the same batch and from the same parents. Behavior experiments were carried out from 11 am - 6 pm.

No animals were excluded from analysis, which was performed using custom-written ImageJ^41^ macros and Matlab codes that allowed automatic tracking following thresholding. Correlation coefficients (Pearson's r) were calculated using the *corrcoef* function in MATLAB (MathWorks). Sample size was chosen based on preliminary experiments, and on previous work with a similar assay^17^. No blinding was done, as a computer performed analysis automatically.

The first 20 seconds of each light / dark cycle was used to calculate initial swimming speed. The last 20-second window was used to compare vertical position. Results of vertical position during migration were presented as (x ± y) representing (mean ± 95% confidence interval (CIs)) to estimate the means of vertical positions of the population. Some data sets rejected the normality hypothesis as determined by the Shapiro-Wilk test. Therefore, all the pairwise comparisons were performed with the Mann-Whitney-U test, a non-parametric test, at a significance level of 0.05. * p < 0.05, ** p < 0.01, *** p < 0.001. The use of the one- or two-tail test is stated in the legend. For multiple comparisons, Kruskal-Wallis tests were performed first, followed by pairwise Mann-Whitney-U tests with the Bonferroni correction. For an experiment with N hypotheses with a desired α = 0.05, the Bonferroni correction were performed on each individual hypothesis at α = 0.05 / N. * p < 0.05 / N, ** p < 0.01 / N, *** p < 0.001 / N. Statistical test and data analysis were performed in MATLAB.

The use of multiple light-dark cycles represents technical replicates; the use of multiple animals represents biological replicates. The occurrence rate is defined as the number of cases where the fish reached the half way point during a climb or dive.

To establish the appropriate intensity of blue light, three different light intensities were tested: 6 μW/cm^2^, 600 μW/cm^2^ and 6000 μW/cm^2^. 600 μW/cm^2^ was used in all subsequent experiments.

### Two-photon calcium imaging

5-14 dpf zebrafish larvae were first anaesthetized in mivacurium and then mounted in 1.5% low-melting-temperature agarose in a glass-bottom dish (Mat Tek) and immersed in E3 water. Fish were imaged on an upright Nikon resonant-scanning two-photon microscope (A1RMP), with a 25x water immersion objective (NA= 1.1) and the laser (Coherent Vision II) tuned to 960 nm. To image different z planes, a piezo drive (Mad City Labs) was used at a step size of 10 μm for light-evoked activity recording and 0.866 μM for the reference brain. The frame rate for whole brain imaging was 7.1 Hz (Figure 2); each stack consisted of 10 frames. For thalamus, habenula and IPN imaging, 4 planes were captured at 1 Hz/stack. For light stimulus, the blue and red light boxes were controlled by a power supply unit (TMS-lite), triggered by a 5 V TTL signal from a National Instruments DAQ board that was controlled by the Nikon Elements software.

Green light could not be used as a stimulus in imaging experiment, due to overlap with the emission spectrum of GCaMP6f.

### Image analysis

For whole brain imaging data, principal component analysis (PCA) was used to reduce dimensions and independent component analysis (ICA) was used to separate independent signals. Matlab codes for these were adapted from a toolbox provided by Mukamel et al^23^. Spatial maps from ICA were used to generate segmented regions of interest (ROIs), and temporal signals from ICA were used to calculate correlation (corr function in Matlab) to the stimuli and between different animals (correlation > 0.5 and p-value <0.05 were used as threshold.).

### Integration of calcium imaging data from multiple animals

5 template signals were generated using square and sawtooth waves: square waves for blue onset and red onset; sawtooth waves for mixed onset, mixed offset and blue offset. The categories and shapes of the template signal were chosen based on observation from all fish data. Blue and red light pulses were delivered randomly. The timing of the templates were determined by the stimulus, and are different for different fishes. For an ICA temporal signal, the correlation coefficients against all the 5 templates were computed and then the maximum was chosen. If the maximal correlation coefficient was larger than 0.5 (threshold chosen empirically), then this ICA signal, as well as its corresponding spatial map, was categorized. Finally, the ICA spatial maps, from different fish but belonging to the same categories, were registered by translation and then averaged.

### Code availability

Macros used in this study will be made available at bitbucket.

### Laser ablation

*Tg(elavl3:GCaMP6f)* larvae were anaesthetized and then mounted in 2% low-melting temperature agarose. Lesions were created with the IR laser tuned to 960 nm and fixed on a single point. Several pulses, each lasting 100 - 500 msec, were used. Lesioning was monitored by time-lapse imaging before and after each pulse, and was judged successful when there was a localized increase in GCaMP6f fluorescence. To further characterize the lesion, a z-stack was collected.

---

**Movie 1. Lesion of the left dorsal neuropil**

A z-stack through the habenula after the neuropil of the dorsal left habenula was lesioned. The cavity induced by the lesion is coloured magenta, while habenula neurons are visible because of the expression of GCaMP3 under the *s1011t* GAL4 driver. Planes are 3 μm apart. Anterior is to the top.

**Movie 2. Lesion of the ventral neuropils**

A z-stack through the habenula after the neuropil of bilateral lesion of the ventral regions of the habenula. The cavity induced by the lesion is coloured magenta, while habenula neurons are visible because of the expression of GCaMP3. Planes are 3 μm apart. Anterior is to the top.

